# A new microfluidic concept for successful *in vitro* culture of mouse embryos

**DOI:** 10.1101/2020.11.10.376160

**Authors:** V. Mancini, P.J McKeegan, A.C. Rutledge, S.G. Codreanu, S.D. Sherrod, J.A. McLean, H.M Picton, V. Pensabene

## Abstract

Innovative techniques for gene editing have enabled accurate animal models of human diseases to be established. In order for these methods to be successfully adopted in the scientific community, the optimization of procedures used for breeding genetically altered mice is required. Among these, the *in vitro* fertilization (IVF) procedure is still suboptimal and the culture methods do not guarantee the development of competent embryos. Critical aspects in traditional *in vitro* embryo culture protocols include the use of mineral oil and the stress induced by repetitive handling of the embryos.

A new microfluidic system was designed to allow for efficient *in vitro* culture of mouse embryos. Harmful fluidic stress and plastic toxicity were excluded by completing the industry gold standard Mouse Embryo Assay. The potential competence of the embryos developed in the device was quantified in terms of blastocyst rate, outgrowth assay, energy substrate metabolism and expression of genes related to implantation potential.

Mass spectrometry analyses identified plastic-related compounds released in medium, and confirmed leaching of low molecular weight species into the culture medium that might be associated to un-crosslinked PDMS.

Finally, these data show the potential for the system to study preimplantation embryo development and to improve human IVF techniques.

## Introduction

Genetically altered (GA) mice are used extensively to study the function and regulation of genes and their role in human development and health. Almost 50% of the total amount of animals used for scientific research are genetically modified animals, the majority of which are mice [1].

GA mouse models provide new insight in the fields of medical research and pharmaceutical industry thanks to the recent introduction of sequencing, genetic engineering technologies (such as CRISPR/Cas9, Microinjection, Embryonic Stem Cell Injection or Nuclear Transfer) and to the use of “humanised” mice, in which a particular mouse gene is replaced by its human counterpart known to be associated with a specific human disease.

Examples of well-known mouse models include important diseases, such as ageing [2], [3] breast cancer [4], [5], metabolic disorders, diabetes, obesity and their use in drug discovery [6], [7].

Improving research animal welfare correlates with robust and high-quality data [8]. The animal facilities for breeding mice use *in vitro* or *in vivo* fertilization techniques to generate and cryopreserve embryos prior to distribution [9], [10]. Many laboratories are currently making significant changes in their breeding protocols and methods to reduce animal suffering, improve throughput and increase birth/pregnancy rates [11]–[13].

The most widely used traditional method for *in vitro* embryo culture in *In Vitro* Fertilization (IVF) relies on 5-100 μl microdrops of specialised embryo culture medium [14] in Petri dishes covered with mineral oil to prevent evaporation (Fig. 1.a). Embryos are transferred to the microdrops and allowed to develop undisturbed in an incubator. When ready to be transferred, developed blastocysts are aspirated and injected in the oviduct or in the uterus by traditional Surgical Embryo Transfer (SET) techniques or by Non-Surgical Embryo Transfer (NSET) [12]. Manipulation of the embryos occurs by manual pipetting, which is labour intensive, time consuming, not repeatable and complicated by the presence of oil. First available in 2009, NSET techniques are now adopted globally and are of similar or improved efficiency to SET [11], [12], [15]. A fundamental requirement for the success of these techniques is the ability to highly control the culture environment of the embryo, producing competent embryos at the required developmental stage. Higher success rates have been reported when culturing embryos to the morula or blastocyst stage prior to transfer, a procedure requiring up to 4 days of *in vitro* culture in the mouse and up to 8 days in other mammalian species [16]–[18].

**Figure 1.**
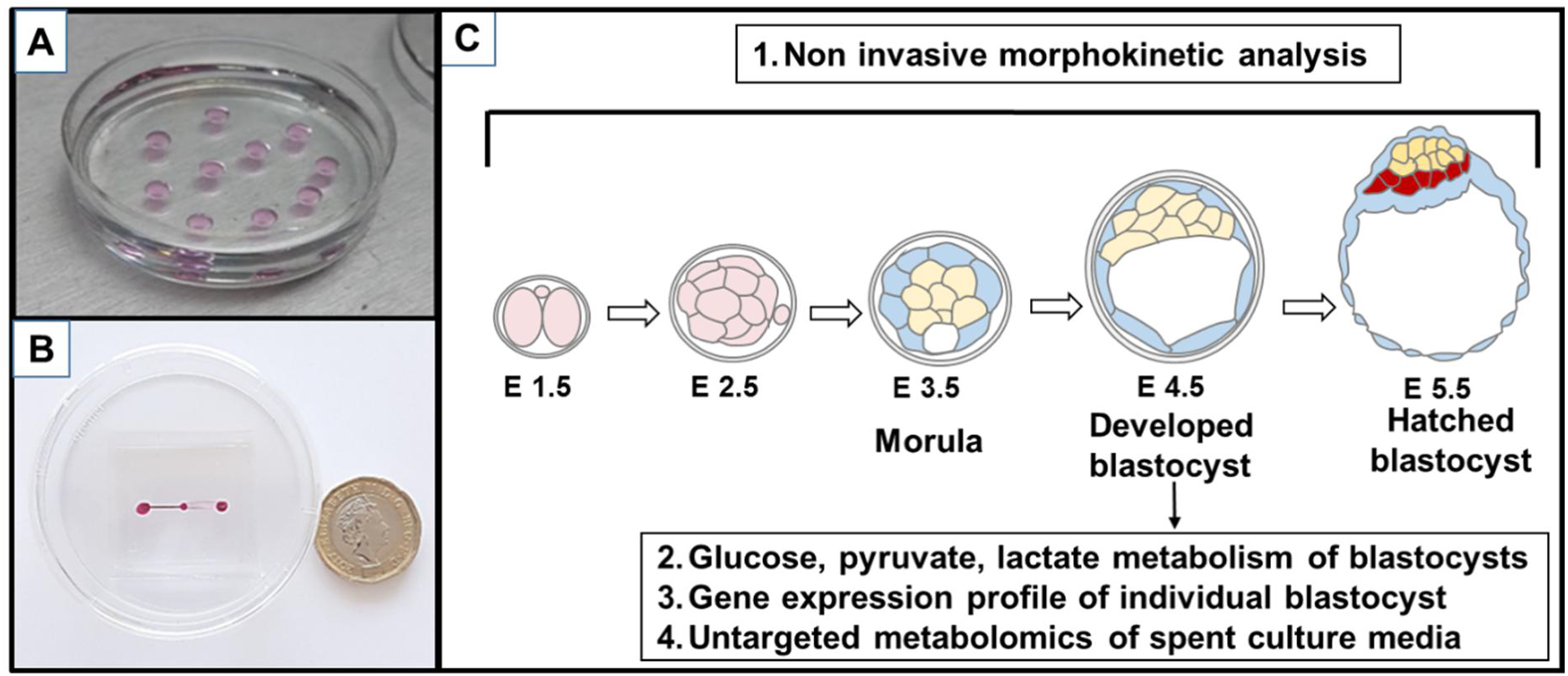
Fig. 1. A) Traditional microdrop culture. B) Image of the fabricated PDMS microfluidic device in a traditional 60 mm petri dish used for embryo culture. C) Schematics of in vitro embryo development in the device. Non-invasive analytical methods can be used to monitor, grade and stage the embryos during culture. Endpoint analyses (genetic and metabolic profiling) can be performed on a single embryo.

Since the 1990’s, microfluidics has been proposed as a new approach to minimize the specialized requirements or ‘art’ required for a successful IVF. In 2002, Hickman utilized a microfluidic device to evaluate the effect of dynamic medium environment through the different stages of embryo development [19]. Later in 2006 and 2010 Cabrera and Heo observed that dynamic perfusion was essential for successful development of 1-cell mouse embryos in a funnel-shaped microfluidic device, positively influencing hatching rate and cell number [20], [21]. While implantation rates and numbers of ongoing pregnancies were similarly improved by the dynamic culture, the new approach was comparable to the traditional microdrop culture approach. More recently, Swain summarized the benefits and limitations of the microfluidic approach for clinical procedures in IVF (these include: maturation, manipulation, culture, cryopreservation and non-invasive quality assessment) [22]. LeGac also presented a comprehensive assessment of the beneficial effect of volume reduction on single and group embryo culture using a microfluidic chamber that supported blastocyst development without altering birth rate [23]. While the interest in the use of these microfluidic systems is still high, extensive scientific analyses to assess safety, consistency and accuracy remain and the long-term impact of culturing embryos in microfluidic devices is not completely understood.

Material toxicity is one of the main concerns, mostly due to the fact that these prototypes are fabricated “in-house” using manufacturing processes difficult to control. The use of polymers that are not medical graded, impure or not completely polymerized affects the safety of the materials in IVF.

Plastics and chemical residues present in manufacturing consumable plastics can not only affect the viability of the embryos, but also trigger long-term adverse effects on the foetal development [24]–[28].

Most of the microfluidic devices proposed in literature for mouse embryo culture are manufactured in poly dimethyl siloxane (PDMS). Data relative to blastocyst rate and birth rate obtained with different devices support the safety of this material. Alternative materials, such as poly methyl methacrylate, polycarbonate, cyclic olefin polymers and copolymers and the most common polystyrene are common thermoplastic materials used in industrial manufacturing of disposable plastic. Most of these are used for microfluidic manufacturing with cheap and fast processes but published evidences do not exclude their long term toxicity [29]. In 2004, Wheeler [30] compared the development of embryos cultured in a microfluidic channel under static conditions, manufactured in silicon/borosilicate and PDMS/borosilicate. In these data, both cases showed improvement of cleavage and blastocysts rates compared to the microdrop culture method. Unfortunately most of these results were not consistent with the Mouse Embryo Assay, which is an industry gold standard for quality control of all lab-ware, media and any other tool that will come in contact with the embryos during the IVF process [31].

Interestingly, the main advantage of the microfluidic approach is the reduction of volumes below the standard microdrop technique. Over the last two decades, extremely low volumes (1.5 to 2 μl) of medium have been used in human ART in the ultra microdrop culture technique [32]. This technique enabled high clinical pregnancy and implantation rates, mostly attributed to the concentration of autocrine and paracrine factors released and available to the embryos in the reduced volume, which are otherwise diluted in the larger volumes necessary in the microdrop technique [33]–[35].

Evaluation of embryo quality indexes provides a thorough understanding of the impact of the microfluidic culture and the reduced volumes on the health and developmental potential of *in vitro*-derived mouse embryos. Morphokinetics (cell cleavage, timing, cell counting, morphology) [36]–[40], energy metabolism [41]–[43], outgrowth assay [44], total cell count, cell allocation and blastocyst rate [45]–[47] represent biomarkers of blastocyst competence that can be evaluated *in vitro* and can be used to optimize the microfluidic technology before moving to *in vivo* testing, thus avoiding unnecessary sacrifice of animals. While some of these methods have been used for validation of microfluidic prototypes, metabolic and genetic signatures of embryos developed in microfluidic devices have never been compared with those of embryos cultured in traditional microdrops.

Mouse preimplantation development is defined by specific patterns in gene expression and cell division, starting from the fertilized egg to final developed blastocyst in approximately 4.5 days. In the last decade, global gene expression analyses have led to the identification of thousands of genes that are expressed and have specific functions during mouse embryo preimplantation development [48], [49].

Analysis of genes involved in trophoblast differentiation (such as caudal type homeobox *Cdx-2* [50], TEA domain family member 4 *Tead4* [51] and E74-like factor 5 *Elf5* [52]) and ICM/epiblast development (such as octamer-binding transcription factor *Oct-4* [53] and SRY (sex determining region Y)-box 2 *Sox2*), or transcriptional repressors such as the methycytosine binding protein genes Mbd [54], can inform about potential effects of the microfluidic environment on embryo development [55],[56].

Finally, difficulties and advantages derived from the adoption of microfluidic systems in standard breeding facilities and correlation of these with standard culture procedures and new less invasive techniques, such as NSET, have not been quantified before, due to inconsistent results and different proposed protocols.

In the current study we developed a novel, oil-free disposable microfluidic device, in which fertilized 1 cell zygote or two-Pro-Nuclei (2PN) stage embryos can expand to blastocyst stage and thus be retrieved for subsequent transfer (Fig. 1.B). Aiming to control and reduce any fluid dynamic shear stress during the handling of the embryos and their injection in the device, the microfluidic design was optimized using Finite Element Modelling (FEM); the model also verified the efficiency of the embryo loading and nutrient diffusion in the system. The potential toxicity of PDMS was assessed by performing the Mouse Embryo Assay (MEA), in which cleavage, blastocyst development and outgrowth rate were used as predictive indexes of implantation. The embryo culture conditions were optimized by studying the effect of group culture on blastocysts development, performing metabolic profiling and determining mitochondrial polarization ratios. Furthermore, the gene expression profiles of blastocysts developed inside the system were compared to those cultured in traditional microdrops using real-time PCR analysis. These data were used to exclude potential genetic alterations induced by the different environment and culture methods. Finally, global untargeted metabolomics was used to identify PDMS-released compounds from culture media extracted from the microfluidic device at different time points (24 h and 5 days).

Based on this extensive evaluation, we demonstrated the ability of the microfluidic system to improve methods for culturing preimplantation embryos, is compatible with other lab equipment (such as optical microscopes, bench-top incubators) and is amenable to traditional handling and analytical procedures (e.g. loading and retrieval with micropipettes, fluorescent staining, and fixation).

#### Significance

##### BENEFIT TO MEDICAL RESEARCH

A more reliable system for embryo culture would allow:

– To increase the adoption of modern gene editing techniques (eg CRISPR/Cas9),
– To avoid the use of cumbersome and potentially toxic, petroleum-derived mineral oil for in vitro culture of embryos.
– To increase the efficiency of generation, phenotyping, archiving and distribution of model mammalian genomes and mouse mutant lines.

##### 3RS BENEFITS

A robust system for development of healthy blastocysts would facilitate:

– Experimental and breeding procedures in mice (about 1.5 million in UK only).
– The reduction of Surgical Embryos Transfers (SET) (excess of 250,000 currently performed globally each year), which are costly, time-consuming, and stressful to mice.

##### BENEFIT FOR ENDANGERED SPECIES AND FOOD LIVESTOCK INDUSTRY

More efficient methods for IVF would increase opportunities:

– To preserve endangered or critically endangered mammalian species (~700).
– To optimize dairy and beef production systems (Blastocyst rates <50% in cattle and sheep).

##### BENEFITS FOR HUMAN IVF

A new device for in vitro embryo culture, validated with a mouse model, could be adapted for human IVF that is:

– Necessary for infertile couples (~10% of the population in reproductive age).
– Suboptimal (< 30% pregnancy rate per IVF cycle).
– Expensive (human IVF cycle >10,000 USD).

## Results

### Microfluidic design is compatible with standard embryo culture methods

The microfluidic device presents two fluidic ports connected to a culture chamber by two microfluidic channels (Fig. 2). The volume of medium in the chamber is 400 nL, thus significantly reduced compared to the traditional 5-50 μL used in microdrop technique and below the volume limit (1.5-2 μL) used in the ultra microdrop method. The inlet and outlet channels are present at different heights and positioned in two different layers (top and bottom): the single inlet channel is 200 μm in height and 250 μm in width. The narrow outlet channels are at lower level and 30 μm tall, thus smaller than the size of a mouse embryo (diameter ~60 μm). These dimensions allow the embryos to be loaded and to reach the chamber, where they cannot flow any further (see Video 1 in Supp. Mat.). Embryos are left undisturbed to grow up to blastocysts (diameter ~100 μm, X days or Y hours) and then aspirated back through the inlet channels to proceed with the transfer.

**Figure 2.**
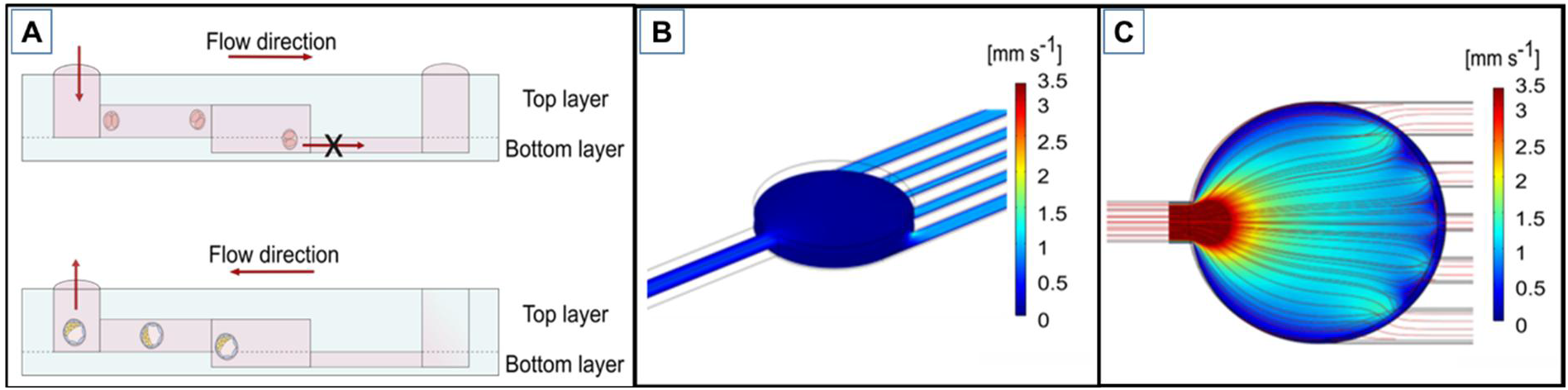
Microfluidic device design. A) Embryos are introduced from the inlet port, guided by capillarity force in the culture chamber through the inlet channel. Being the embryos diameter bigger than the outlet channels, they remain trapped in the chamber. Once developed embryos are aspirated from the inlet port. B) The Finite Element Model of the average velocity shows an increase in the lateral channels compared to the main chamber owing to their small dimensions. C) Fluid flow analysis of velocity magnitude surface plot shows the velocity field generated in the fluidic systems.

The final device, measuring 4 by 4 cm, can be placed in an IVF 60 mm Petri dish, sterilized and used in a MINC™ Mini incubator (Figure 1.B).

### Minimal stress or damage observed by flow and shear stress during loading and through the development process in the microfluidic device

The design of the device allows for the capture of the embryos in the middle chamber owing to the increased hydraulic resistance encountered when reaching the chamber (Fig. 2). Furthermore, the wall shear stress (*WSS*) generated within the channels during loading and retrieval reaches maximum values of 0.17 dyn cm^−2^, which is 7 times lower than values presented in literature (1.2 dyn cm^−2^ shear stress caused lethality within 12 h for E3.5 blastocysts [57] - Supp. Material Fig.S1).

The wide culture chamber and its consequent lower hydraulic resistance [58] prevent embryos to be stressed by lethal WSS during the loading process.

During fluid loading, the velocity profile in the inlet channel (hydraulic resistance in the order of 10^10^ Pa s m^−1^) is significantly reduced in the culture chamber (where an evenly blue the outlet channel due to an increase in hydraulic resistance (∝ 10^12-13^ Pa s m^−1^). The inlet and outlet channels have similar fluid velocity profiles. The device was designed so that the total width of the outlet channels is comparable to the whole chamber width, thus the velocity profiles within the culture chamber are optimized for embryo growth along the cross-sectional direction (Supp. Material Fig. S2).

An additional design criterion was to favour the spreading of the embryos in the whole chamber and to ensure a homogeneous perfusion of medium in the chamber. As shown in Fig. 2.C and in Supp. Material Fig. S2, a wide velocity profile of the optimized device design was obtained within the culture chamber along the cross-sectional direction when the device included multiple channels disposed parallel to the inlet channel. Real flow characterization performed by injecting fluorescein in the microfluidic chamber, showed that the green dye diffused and reaches equilibrium in 8 sec, as reported in Suppl. Materials Fig.S2.

After thawing and washing with fresh medium, embryos are loaded in the device using a flexi pipette (EZ-Grip, 145 μm); these pipettes are traditionally used for embryo culture and has a tip size compatible with the inlet port of the device. The embryos are manually injected into the inlet port and owing to fluid movement they safely move through the inlet channels and reach the central chamber (Suppl. Material Movie 1).

The microfluidic device was designed to minimize the clustering of embryos, even when cultured in larger groups (20, 30 and 40 embryos).

### Experiment 1. Blastocysts rate, hatching rates and outgrowth are not affected by the microfluidic environment

Assessment of morphology at distinct time points is regularly used for evaluating embryos’ quality. Early cleavage, occurring on average at 24h after pronuclear fusion, has been correlated with embryo quality together with blastocyst rate and hatching rate both in mice and in human. Cryopreserved zygotes were thawed and cultured in KSOM medium in groups of 10 in standard 10 μL drops and in microfluidic devices. As summarized in Fig. 3, embryo development during microfluidic device culture was not altered when compared to traditional culture. Specifically, no significant differences were observed in cleavage rate, blastocyst rate or hatching rate.

**Figure 3:**
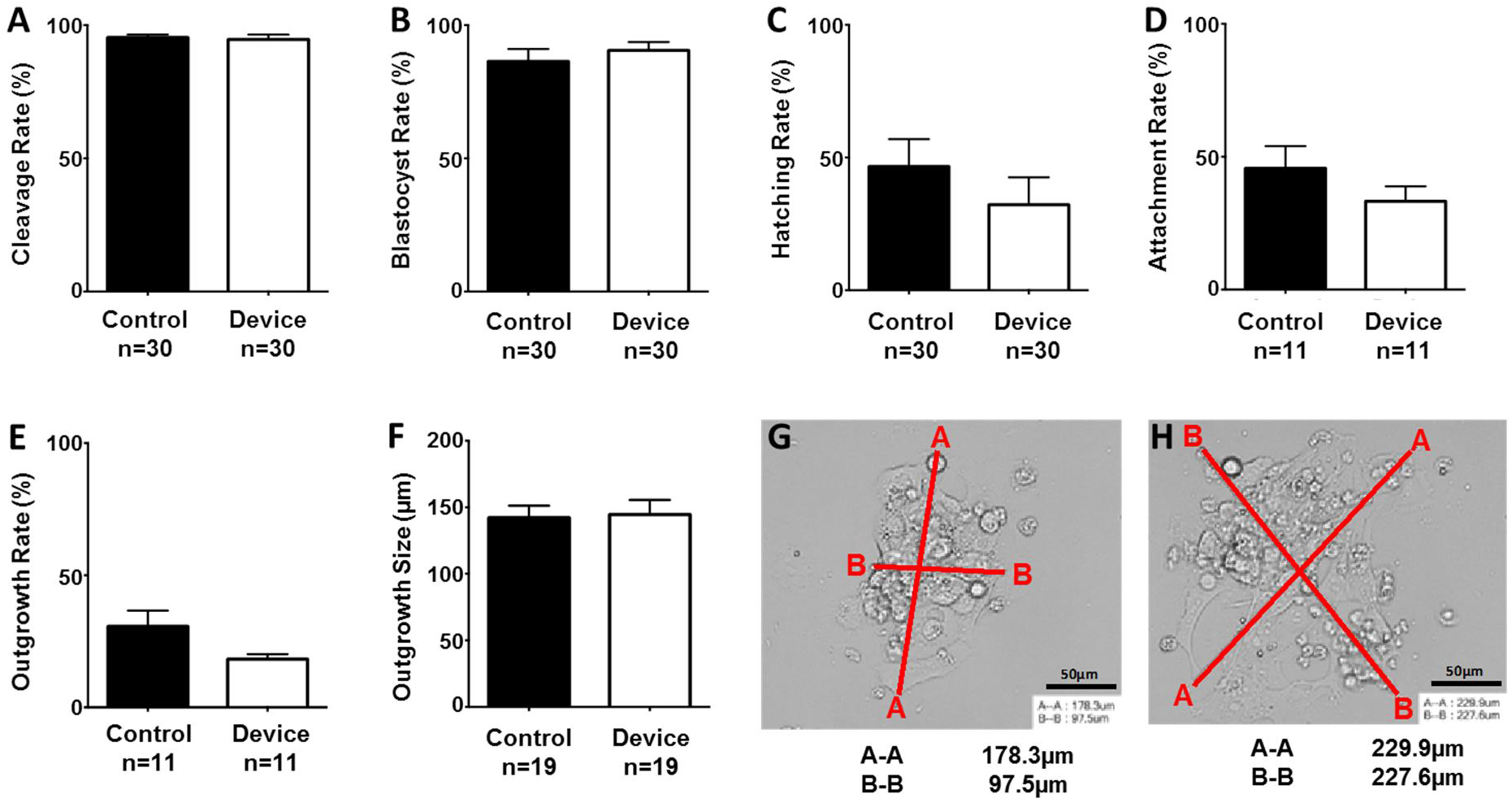
Comparison of preimplantation embryo development. A) Cleavage rates (94.62 ± 1.7%, n=30 vs control 94.84 ± 1.08%, n=30, p=0.48). B) Blastocyst rates (out of total embryos cultured) (90.48 ± 3.2%, n=30 vs control 86.40 ± 4.6%, n=30, p=0.6), C) Hatching rates (out of total blastocysts) (32.18 ± 10.35%, n=30 vs control 46.61 ± 10.28%, n=30, p=0.32), D) Attachment rates (out of total blastocysts) (33.22 ± 5.6%, n=11 vs control 45.64 ± 8.4%, n=11, p=0.35), E) Outgrowth rates (out of total blastocysts) (18.2 ± 1.9%, n=11 vs control 30.6 ± 6.0%, n=11, p=0.08), F) Mean outgrowth size (μm) (144.5 ± 11.0 μm, n=19 vs control 142.1 ± 9.0 μm, n=19, p=0.72). G-H) Representative images of blastocyst outgrowth by embryos cultured in microdrops and microfluidic devices respectively: red lines indicate measurement bars created in RI viewer (A-A and B-B). Scale bar 50 μm.

Blastocyst adhesion competence *in vitro* was quantified by transferring embryos into fibronectin-coated plates for analysis of attachment and outgrowth formation (Fig. 3). This assay mimics the natural mechanism of adhesion of trophoblasts on the blastocysts to the endometrium which is regulated by interaction between the integrins naturally expressed on endometrial cells and on the apical surface of competent blastocysts [59–60]. By assessing the extension and adhesion of the hatched blastocysts it is possible to evaluate their developmental potential. The blastocyst attachment data were similar between microfluidic device (33.22 ± 5.6%, n=11) and microdrop-cultured embryos (45.64 ± 8.4%, n=11); these results were not statistically significant different from each other (p=0.35). Some blastocysts did attach without hatching. Outgrowth rate, defined as percentage of blastocysts forming outgrowths (18.2 ± 1.9%, n=11 vs control 30.6 ± 6.0%, n=11, p=0.10) and mean diameter of outgrowths formed were also not statistically different (144.5 ± 11.0μm, n=19 vs control 142.1 ± 9.0μm, n=19, p=0.72).

Cell number in a preimplantation embryo is directly correlated to the embryo health, developmental potential and ability for cell cycle progression. Total cell counts and cell allocation ratios in Day 7 blastocysts was carried out with an antibody-free differential staining method (Fig.4C) [72]. Total cell numbers were similar between controls and devices (Fig. 4A). There were also no significant differences between cell allocation to trophectoderm or inner cell mass lineages as expressed as percentage trophectoderm of total cells (Fig. 4B). The similar allocation and formation of the extraembryonic cell lineage – trophectoderm and the inner cell mass (ICM) supports the negligible effect of the microfluidic confinement on embryo potency.

**Figure 4.**
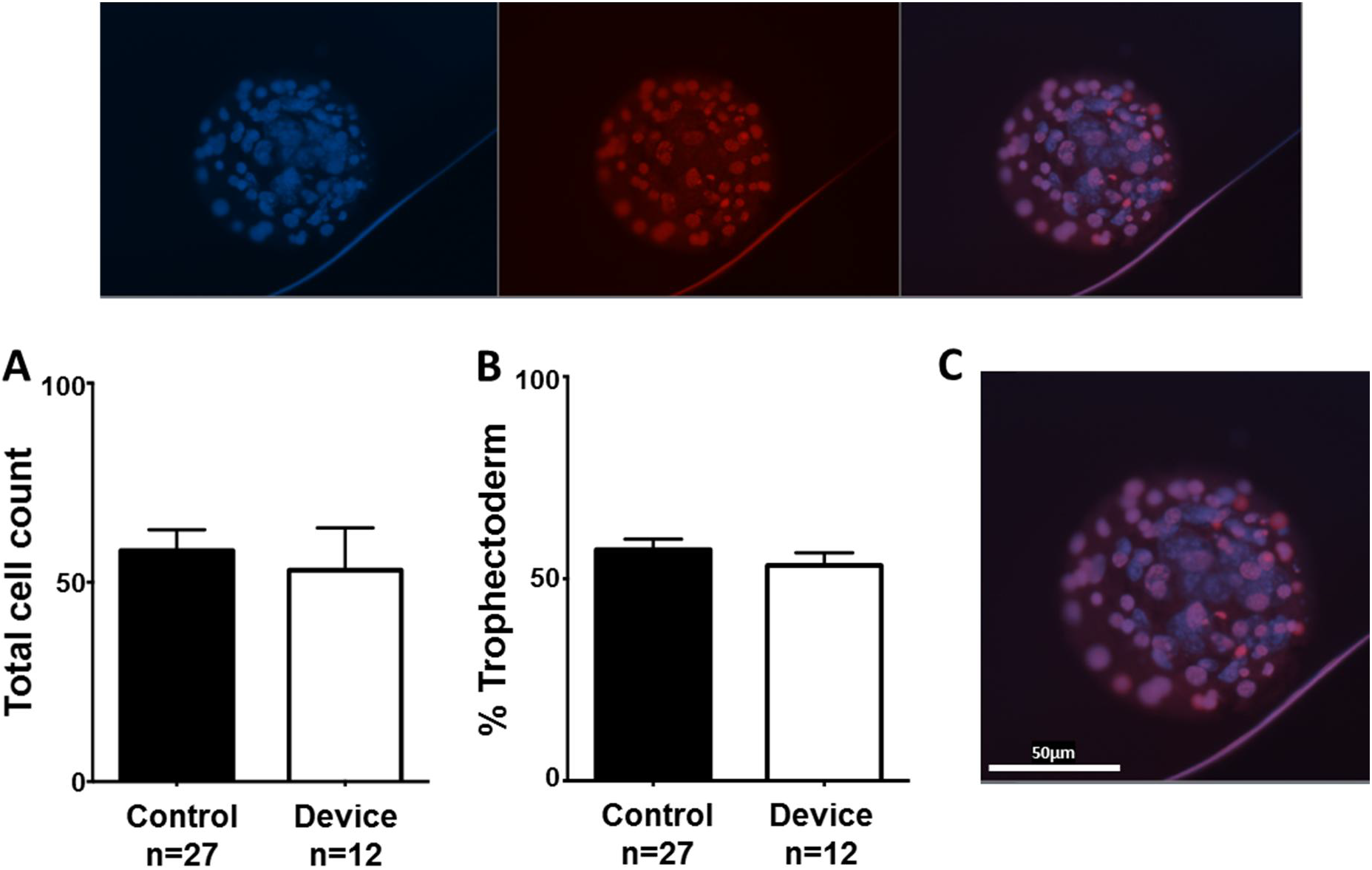
A) Total cell count 53±10.6 cells, n=12, p=0.6 vs control 58± 5.2 cells, n=27. B) Cell allocation ratio (% Trophectoderm) 53.3±3.2%TE, n=12, p=0.4 vs control 57.2±2.6%TE, n=27, p=0.4. C) Representative conventional epifluorescent image of blastocyst within the device stained with Hoechst 3342 and Propidium Iodide and imaged in the 460nm (blue) and 560nm (red) channels respectively. Scale bar 50 μm.

### Experiment 2. Embryo group size limits development and metabolic activity in the microfluidic chamber

The number of embryos cultured in a single microdrop and the volume of the microdrop are two variables that affect embryo development during culture. Different protocols have been compared to identify an optimal range of embryo density, in terms of μL of medium available per embryo. In a microfluidic system, the embryos are confined in a different environment, specifically a 400 nL static chamber is linked through two side channels and allows two 10 μL drops of medium to be available to provide nutrients during ~3 days of culture. It is extremely important to evaluate how the different fluidic environment can support embryo growth and provide sufficient nutrients.

When comparing groups of 10, 20, 30 and 40 embryos, the overall blastocyst rate was not significantly different between groups of different size (p=0.25) (Fig. 5). It was highest in the device 10x group, and decreased with the size of the group, from ~90% in the 10x group down to ~56% for the larger group of 40 embryos. Groups of 10 embryos very consistently gave 80-90% blastocyst rates, while larger group sizes were very variable between replicate cultures. However, hatching rate in devices significantly decreased with increased group size (10x group: 30±4% compared to 40x group: 2.2±2%, n=4, p=0.02). This suggests that substrate competition outweighs increased paracrine effects in larger group sizes.

**Figure 5.**
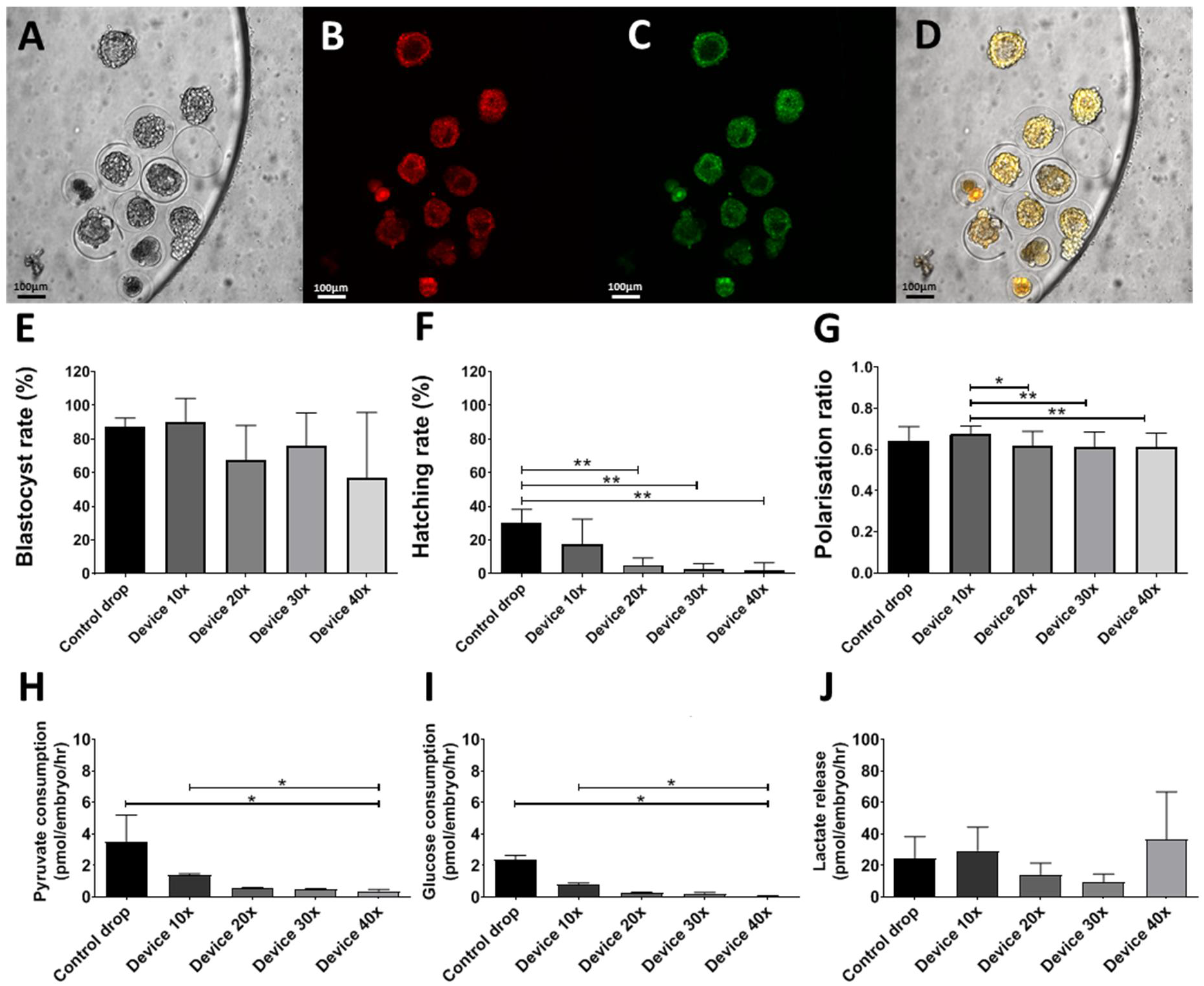
Loading capacity experiment. A) Brightfield, B-D) Analysis of mitochondrial polarisation ratio. Representative conventional epifluorescent images of blastocysts within the device stained with the potentiometric mitochondria-specific stain JC-10 and imaged in the rhodamine isothiocyanate (RITC, B) and fluorescein isothiocyanate (FITC, C) channels respectively. Scale bar 100 μm. E) Blastocyst rate (n=4) F) Hatching rate (n=4) G). Quantified blastocyst polarisation ratio data (n=4) shows higher value when culturing groups of 10 embryos inside the device. H-J) Energy substrate turnover (n=4).

Development rates are a widely used key marker of ongoing developmental competence. However, to examine embryo competence in richer detail, metabolic parameters were investigated. Blastocysts were labelled and imaged directly within the devices using the ratiometric mitochondrial dye JC-10 (Fig. 5) to evaluate changes in mitochondrial membrane potential. JC-10 accumulates in the matrix of polarised mitochondria, forming J-aggregates with punctate red fluorescent signal. In areas of reduced mitochondrial polarisation, the dye tends to remain in monomeric form with diffuse, green signal. Mitochondrial polarisation thus indicates overall mitochondrial polarisation in mammalian oocytes and embryos [61–62]. Control blastocysts cultured in microdrops did not have a significantly different mitochondrial polarisation ratio (0.64±0.02, p>0.92) compared to device-cultured embryos, regardless of the groups size. However, following device culture, blastocyst polarisation ratio was significantly higher in 10x groups (0.67±0.04, n=27) compared to 20x groups (0.62±0.01, n=42, p=0.01) 30x groups (0.61±0.07, n=71, p=0.0005) and 40x groups (0.61±0.7, n=66, p=0.001). Overall blastocyst mitochondrial polarisation ratio correlated strongly with blastocyst rate across all groups (p=0.01), confirming the viable status of the embryos.

Control embryos had metabolic profiles typical of microdrop-cultured murine blastocysts [63–42]. Device-cultured embryo pyruvate and glucose consumption decreased with increasing group size. Embryos cultured in groups of 40 had significantly reduced pyruvate (0.37±0.1 pmol/embryo/hr) and glucose consumption (0.05±0.03 pmol/embryo/hr, n=3) than groups of 10 (1.4±0.08 pmol/embryo/hr, and 0.8±0.08 pmol/embryo/hr, n=3, respectively p=0.02). Pyruvate is the preferred energy substrate during early cleavage, while glucose consumption is low during early cleavage but tends to increase greatly with increased ATP generation through oxidative phosphorylation at the blastocyst stage [42]. The present data suggests increased competition for these substrates within the more concentrated population of 40 embryos in comparison to groups of 10 in either devices of control microdrops. Device embryos were more quiescent overall, with reduced variation between culture groups. Embryos with metabolic profiles in the intermediate or *lagom* range and with a tighter distribution may be the most developmentally viable, due to undergoing less metabolic stress [42]. The minimal, physiologically accurate volume of available medium in device culture may encourage embryos towards this moderate metabolic profile and improve embryo developmental potential.

### Experiment 3. Microfluidic confinement does not affect development and implantation related gene expression

Using Real-time PCR (qPCR) of single blastocysts, the expression of 53 genes involved in blastocyst development and cell differentiation (Fig. 6A) was analyzed. In particular, differential expression of marker genes involved in trophectoderm differentiation (*Sbno1* [64], *Klf5* [65], *Cdx2* [66], *Tead4* [51], *Gata* [66], *Elf5* [52], *Krt18* [50]) or ICM/epiblast development (*Stat3* [67], *Nanog* [50], *Pou5f1* [68], *Sall4* [67], *Gata6* [69]) was measured (Fig. 6B). No statistically significant differences were observed in these genes when blastocysts developed in the microfluidic device were compared to those cultured in traditional microdrops. These results confirm that the device does not affect gene expression of mouse blastocysts, and therefore suggest that the microfluidic device culture environment is not harmful to mouse embryos.

**Figure 6.**
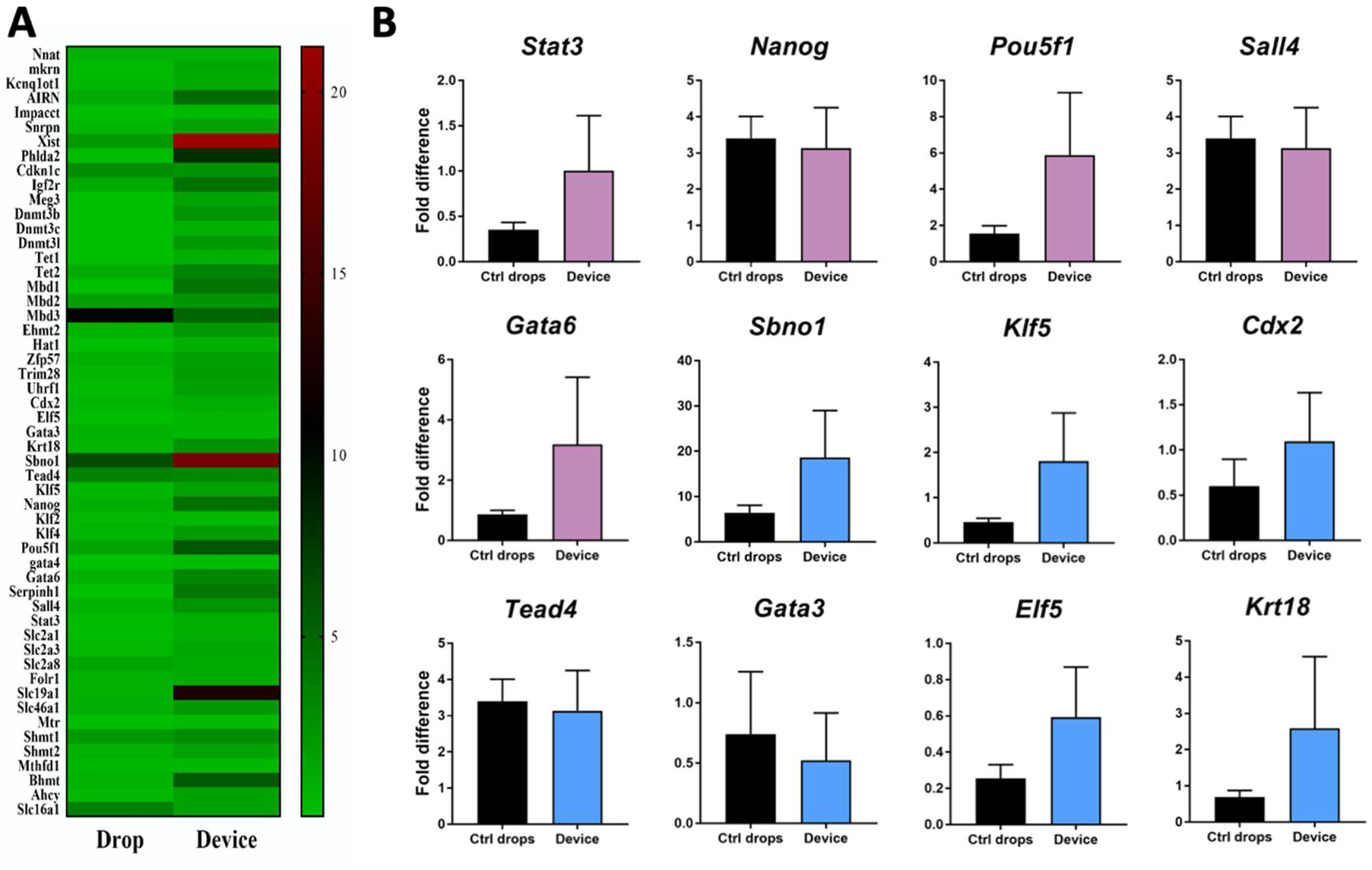
(A) Heatmap representing gene expression of mouse blastocysts cultured in the device compared to microdrop culture. Scale: red indicates high expression and green is low expression. (B) Relative mRNA expression of selected genes (purple: markers of ICM/epiblast differentiation; blue: markers of trophectoderm differentiation) in blastocysts (n=10) cultured in the device, compared with control (n=10). Data are presented as mean ± SEM.

### Experiment 4. PDMS alters the medium composition but does not induce drastic changes in the embryo metabolism

Global, untargeted liquid chromatography tandem mass spectrometry (LC-MS/MS) analyses were performed to compare spent medium (KSOM) on days 4 or 5 of embryo culture between each experimental group (KSOM, KSOM + microdrop, and KSOM + device). The metabolite composition of spent KSOM collected from the microfluidic device was compared with that of spent medium collected from control microdrops, both in the presence or absence of embryos (n= 3 for each group). The Principal Component Analysis (Fig. 7A) and heatmap (Fig. 7B and Fig. S3) show distinct clustering of the experimental groups. These global views revealed that while the majority of the detected metabolites were stable in abundance, there was a modest subset of metabolites in culture medium with unique abundance profiles between device and microdrop technique.

**Figure 7.**
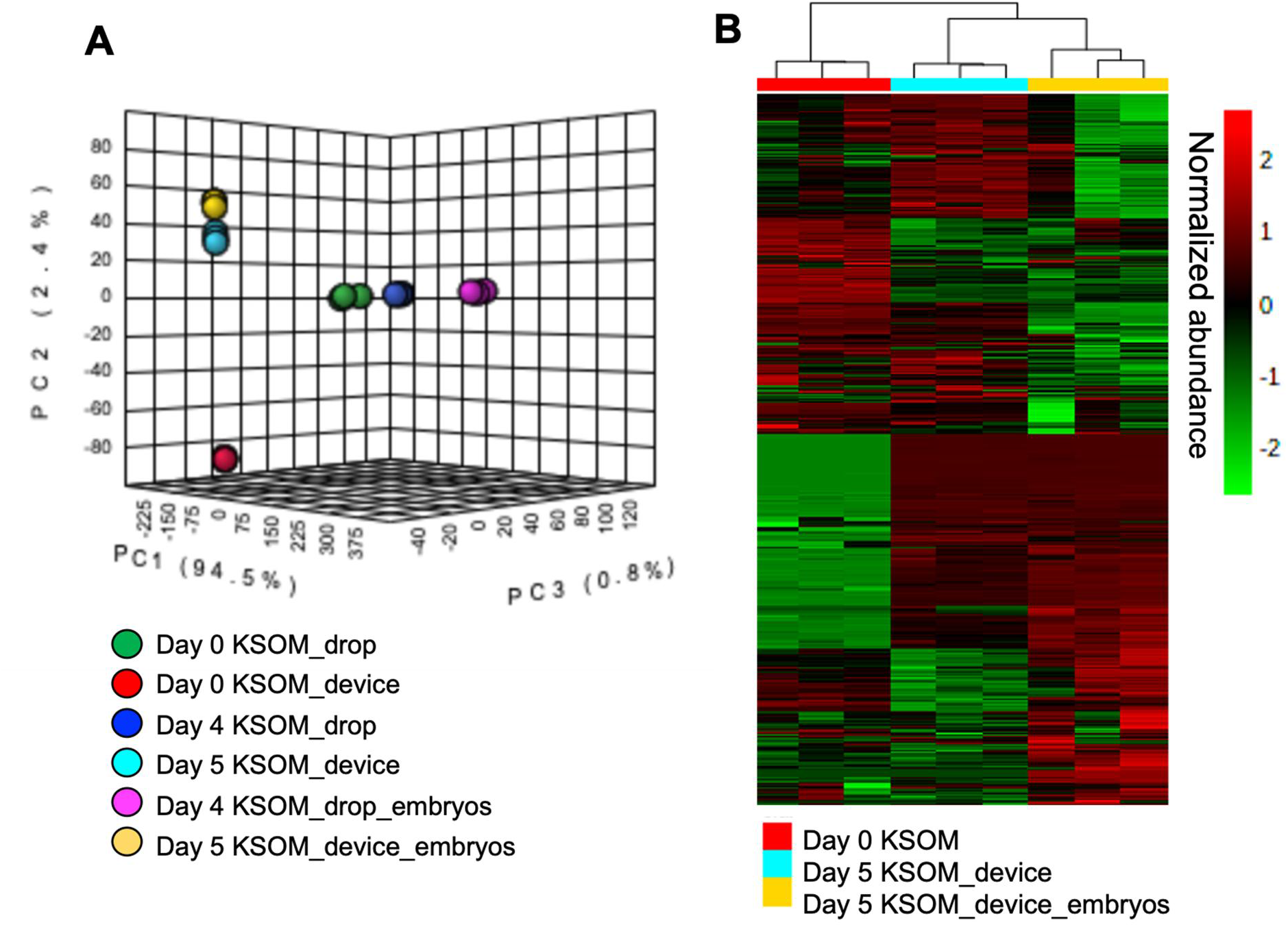
Principal Component Analysis (PCA) and heatmap visualization of global, untargeted mass spectrometry. (A) PCA plot of the LC-MS/MS data of medium samples from devices after 4 days in microdrops or 5 days in devices and control KSOM (n=3 per experimental group). (B) Heatmap analysis of media samples collected from the microfluidic device with and without embryos. Sample replicates are visualized in columns column based on hierarchical clustering, with metabolites presented on individual rows. Species are colored based on normalized abundance from red (high) to green (low).

Second-order meta-analysis of the individual pairwise comparisons (Fig. 8) allows prioritization of endogenous or xenobiotic compounds (specific to PDMS, PS or mineral oil exposure) that were altered in abundance by embryo culture method (device vs. microdrops). The Venn diagram shows shared and unique compounds for specific comparisons. A comparison of the compounds identified in spent media from devices (“PDMS-media”) or spent media from control microdrops on standard Polystyrene dish (“PS-media”) without the presence of embryos revealed a total of 547 compounds (Fig. 8A). Using meta-analysis, 48 compounds were common to the two groups (device vs. microdrop), whereas 387 were unique to the PDMS-media group and 64 to the PS-media group. Compounds unique to the device group, represent species (xenobiotics) directly associated with PDMS use. These data show 339 xenobiotic compounds were released by PDMS into the culture medium and 48 compounds were absorbed by or adsorbed onto the PDMS from the culture medium. Included in the 339 species released into the culture medium, plasticizers such as butyl lactate, dimethyl sulfoxide, ethanol, 2-[2-(2-butoxyethoxy)ethoxy]-, N,N-dimethylformamide, necatorine, pentaethylene glycol, triethylene glycol and tripropylene glycol were detected. The effects of these potentially toxic compounds on embryo development must to be further investigated. The remaining detected compounds indicate breakdown products of media components produced by media degradation after 5 days of incubation at 37 °C. These include peptides, amino acids and other small endogenous molecules (e.g., l-glutamine, l-tryptophan, n-phenylacetylglutamic acid, pyridoxamine, dihydrolipoamide).

**Figure 8.**
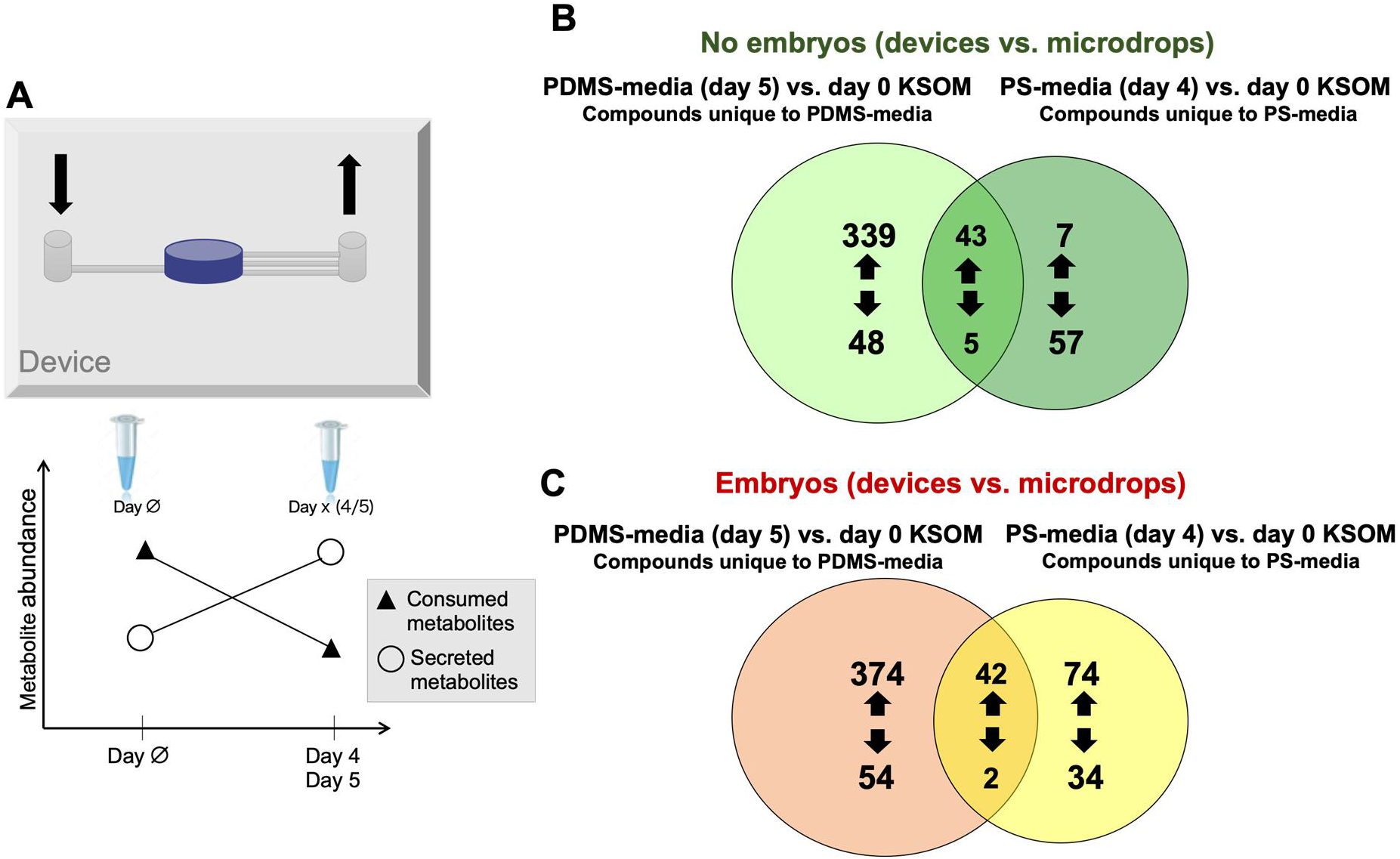
Venn diagram of dysregulated compounds in PDMS-media and PS-media compared to control (day 0 KSOM). (A) Schematic of the device with arrows indicating inlet (↓) and outlet (↑) ports (top). Changes in metabolite abundance in samples collected from microdrops (day 4) or devices (day 5) when compared to control. Increased and decreased compounds represent, respectively, released and consumed metabolites (bottom). (B) Comparison of dysregulated compounds in day 5 PDMS-media and day 4 PS-media without embryos. (C) Comparison of dysregulated compounds in day 5 embryo culture PDMS-media and day 4 embryo culture PS-media.

Similarly, among the 48 species that were significantly decreased in spent media from the device (Fig. 8A), numerous biological compounds were identified that were sequestered by PDMS from the culture medium, these include: amino acids and dipeptides (e.g., isoleucyl-isoleucine, isoleucyl-leucine, isoleucyl-phenylalanine, n-acetyl-l-methionine, and valyl-leucine).

Among the species that were significantly decreased (57 compounds) in spent media from microdrop culture, peptides and amino acids, such as L-tyrosine, aspartylphenylalanine and phenylacetylglycine were identified. These data suggest that these molecules were absorbed or transformed by the plastic or mineral oil. The remaining detected compounds represent breakdown products of culture media that degraded over time.

Other organic species appeared downregulated in PS-media and PDMS media suggesting a segregation of compounds from the medium both into the elastomer as well as in the plastic or the mineral oil used to prevent media evaporation. These molecules include tryptophol [xylosyl-(1->6)-glucoside], muramic acid, [3,5-dihydroxy-2-(hydroxymethyl)-6-[3,5,7-trihydroxy-2-(2,4,5-trihydroxyphenyl)-3,4-dihydro-2H-1-benzopyran-6-yl]oxan-4-yl]oxidanesulfonic, and 2-methoxyestrone 3-glucuronide.

Subsequently, second-order meta-analysis of spent media collected with embryos present in the device and in the microdrop method was performed. Specifically, embryo cultured spent media from devices (day 5) were compared to embryo present spent media from microdrops (control, day 4) (Fig.8B). These analyses allow for us to determine metabolites and/or xenobiotics present in spent media after embryo culture (i.e., produced or consumed by the embryo during culture) as well as xenobiotics unique to device culture (i.e., associated with the fabrication process or released/absorbed by PDMS) or to control microdrop culture (i.e., associated with PS or the mineral oil used to prevent medium evaporation). The Venn diagram shows 428 species to be only present in embryo cultured PDMS-media, 108 compounds were only observed in embryo culture PS-media and 44 species were present in both embryo culture PDMS-media and embryo culture PS-media (Fig. 8B). Notably, among these 44 common metabolites, the relative abundance of 42 of these compounds were statistically significantly increased (p<0.05 and fold change ratios > 2) in both the embryo culture PDMS-media and the embryo culture PS-media when compared to control, these identified compounds include metabolic markers of pre-implantation embryo development, such as pyroglutamic acid, 5’-methylthioadenosine [70], hypoxanthine [71–72], cytosine, n-acetyl-l-methionine, and phenylacetylglycine. Pre-implantation embryo development compounds represent metabolites produced by embryos in both culture conditions. Similarly, two compounds significantly decreased in spent embryo culture media compared to control (i.e., muramic acid and meticillin) are known to be consumed by embryos from the culture medium.

These results allow to conlcude that expected metabolomic changes can be detected by mass spectrometric analysis in samples of media collected during embryo culture, and these can be correlated with organic and inorganic compounds available to the embryo during its development.

## Discussion

The presented microfluidic device, in accordance with similar platforms developed in the last decade for IVF, resulted a simple and user-friendly system for culturing murine embryos and enabled the use of several, non-invasive techniques for assessing embryo quality.

The individual microfluidic compartments of the device were designed so that user friendly efficient, reproducible loading and recovery/ retrieval of the embryos was achieved. The procedure for culture preparation was facilitated by the absence of oil and by the compatibility of the device with bench top incubators and optical microscopes.

Embryo development was similar to the widely adopted microdrop method, with both cleavage and blastocyst rates exceeding 90% in all the conditions. Taken together, these data indicate that the novel microfluidic device presented here is embryo-compatible and can successfully be used to support embryo development to the blastocyst stage without the need for media changes or oil overlay.

The analysis of gene expression of single blastocysts fully developed in the microfluidic device provided a clear assessment of the influence of the microfluidic confinement on the embryo quality. The selection of genes correlated to the preimplantation and endometrial receptivity provides evidence to support limited alterations of the embryos during the 5 days of culture. The combination of gene and metabolite data analyses are complementary, metabolomics data allows us to identify at the molecular level metabolites released by the embryo or consumed by the embryo during development while genomic data allows us to follow changes of embryo development at the gene and protein level. Interestingly, the cultured medium is altered significantly at day 0, in accordance with previous work on PDMS absorption [73] and the release of xenobiotic compounds in the medium is present in a few hours. These compounds can be ascribed to the unstable hydrophilic properties of the PDMS over time. These properties influence leaching and sequestration of small molecules by the elastomer [74]. As revealed by the culture medium analysis, specific compounds can be found uniquely in the samples derived from the microfluidic system. In these data, numerous uncrossed-linked components of the PDMS were identified. Second-order meta-analysis allowed for biologically important metabolite changes to be observed in the culture medium throughout embryo development. Identification and quantification of metabolite and xenobiotic compounds is not completed at this stage, however these data show that the presence of the embryo in culture altered the composition of the medium in the device and in the microdrop method. These data also show that the number of common compounds in the different culture settings (device vs. microdrops) were changed. However the embryos developed successfully in the PDMS device, without significant alterations. The stability of PDMS may represent a challenge in the field of microfluidics and lab-on-chip research, however alternative plastics for manufacturing of the device could be used for this novel system. The methods used for evaluating the embryo development in this work should be performed at minimum for industrial manufacturing plastic. Importantly, from the morphokinetic analysis, these unexpected xenobiotic compounds observed in the medium did not induce significant alteration of the embryo development at different stages.

Oxygen availability is also a key requirement for embryo development [75]. In this work, we assessed the effect of the confinement of the embryos in a close compartment, completely surrounded by PDMS, which is permeable to gas [76–77]. Oxygen tension of 5% for embryo culture has been widely adopted in both animal and clinical human IVF laboratories and considered a more physiological concentration that can boosts blastocyst development with no detectable adverse effects [78–82]. While it is true that the diffusion of oxygen through the PDMS could be modelled and compared to that through mineral oil and media, we did not observed detrimental effects on the embryo development due to a different oxygen dynamic. The microfluidic device allows to collect single blastocysts and to analyse their specific genetic profile by real time qPCR. In this study the relative expression of 12 specific genes involved in blastocyst development and cell differentiation, trophoblast and epiblast development, aimed to exclude any genetic alteration induced by the microfluidic environment. In order to use the same technique and to correlate those specific genes to the blastocyst competence, a more consistent breeding protocol would be required, with standardization of fertilization method, sperm and egg donor and including different mouse strains.

The LC-MS/MS analysis of spent media revealed changes in the media composition, with sequestration of molecules and different consumption and release of compounds by the embryo in the microfluidic device and in the standard microdrop method. These data improve our understanding of the embryo metabolic activity and confirm the unstable characteristics of the prototyping elastomer. A more detailed analysis could be set up by collecting samples every 2 hours to correlate altered media composition with abnormal embryo developmental rates to morula or blastocyst stage.

Future analyses of the medium composition will be used to search for specific toxins, such as peroxides derived from the mineral oil, zinc, and other unknown contaminants that may be released into the medium from the oil or the plastic.

Finally, these data show the feasibility of performing both PCR and metabolomics analyses from single stage matched blastocysts. The robust gene expression data generated from the microfluidic device shows the capability of profiling of preimplantation embryos in a microfluidic environment and confirms the impact microfluidic technology has in fundamental research in assistive reproductive technology.

Further investigation will explore the effects of device culture on embryo implantation and live birth rates. Embryo transfers in mice and assessment of birth rates will confirm the competence of embryos cultured in the microfluidic device and will allow to quantify time and costs saving introduced by this protocol. Preliminary data gathered so far on embryo transfer with BL6 mice [83] has shown improved birth rates (53%) when transferring embryos cultured for 48hrs in the devices with surgical procedures. When combined with NSET, the microfluidic culture ensured successful implantation of *in vitro* matured blastocysts (success rate >25%). Further trials are now planned to evaluate the improvement in terms of birth rate, including different strains of the recipient mice, cryopreserved or fresh embryos and using specific protocols for embryo transfer.

This novel microfluidic technology eliminates the current standard use of potentially toxic mineral oil for embryo culture, thus reducing toxicity, failure, and costs for each culture set up. The loading and retrieval are significantly simplified compared to the microdrop technique, allowing to pipette an entire group of ten embryos in a single step. This represents an additional value in high throughout facilities to maximize the costs and the success of the culture. More importantly this new method also favours the adoption of NSET. Based on our first trials [83], by using the microfluidic dish in combination with NSET [84], animal breeding labs will eliminate the costs related to bench and animal preparation for surgery (i.e. setting up, packaging, sterilisation and cleaning of surgical tools, health monitoring, anaesthetics, analgesics, etc.), will reduce the times required for embryo transfer from one hour to 15 min, and for training lab technician, from four months to one month.

Considering human IVF technique, the microfluidic device adaptable for the culture of human embryos. The technology and the fabrication allow to change the design of the device to accommodate a different and limited number of human embryos and to make it compatible with clinical procedures. To this end, more data are required to move to preclinical trials, in order to prove the effective improvement of the embryo development and exclude any long term, epigenetic effect of the materials, as well as any interference with the current protocols and standards in IVF clinical settings.

## Materials and methods

### Microfluidic device flow and shear stress analysis

COMSOL Multiphysics 5.2a was used to evaluate and compare flow rate, velocity field and predict shear stress as function of microfluidic device geometry.

To generate 3D model, the microfluidic design was first created by using computer-aided design software (Autodesk AutoCAD 2017). The design geometry was then imported into COMSOL library. The fluid inside the device was simulated as an incompressible, homogeneous, Newtonian fluid with density (*ρ*=1000 kg m^−3^) and viscosity (*μ*=1×10^−3^ Pa s) [80].

Flow was created by manually loading a solution of 4.8 μm fluorescent polystyrene beads using an embryo handling Flexipette with 170μm tip (EZ-Grip, RI). The maximal velocity in the inlet channel was estimated to be respectively about 0.4 mm s^−1^(as average of 10 measurements) which corresponds to a flow rate equal to 1.17 μl min^−1^ (Re: 0.087). The devices showed a widely lower flow rate compared to the theoretically derived harmful value previously calculated (13.91 μl min^−1^).

The estimated inlet velocity was applied to a COMSOL model to predict the shear stress the device during embryo loading (Suppl. mat., Figure S1). Critical values of shear stress (~3.5 dyn cm^−2^) were only found on the edges at the interface between culture chamber and outlet channels, areas that embryos cannot reach because of the narrow outlet channels section.

Once velocity has been assessed, to prove the computational model results and to better characterize flow profile inside the microfluidic chambers a solution of fluorescein (0.05 mg ml^−1^) was flowed into the chamber previously filled with water. As predict from the computational analysis, the microfluidic device gives a flow profile as wide as the chamber width (Fig.3.B, Supplementary material, Figure S2). Considering the optimal diffusion of nutrient factors within the chamber (e.g. fluorescein) that this can provide, and the transient mechanical stimulation exerted on the embryos, these results suggest that embryos homogeneously spread in the chamber and receive equal amount of nutrient. These results are promising considering previous data that demonstrated that placing and maintaining of embryos at a specific reciprocal distance of approximately 165 μm enhances embryo growth by favouring paracrine signalling between embryos, and avoiding detrimental accumulation of secreted products in the medium surrounding them [85].

### Microfluidic device fabrication and preparation for culture

Microfluidic devices were fabricated in polydimethylsiloxane (PDMS, Sylgard® 184, Down Corning, MI, USA). This microfluidic structure was obtained by bonding together two layers of PDMS (Figure 1. B), where microchambers and microchannels are defined by standard soft lithography techniques. Once assembled, the devices were immediately filled with sterile tissue culture water and stored closed at 4°C to preserve hydrophilicity. Before embryo culture, devices were sterilized by exposure to UV light (254 nm wavelength for 30 minutes). These were thus prepared by drawing 10 μl KSOMaa (Millipore, UK) from the inlet through the channel and chamber. Devices were then primed by overnight incubation with 10 μl KSOMaa media drops added to the inlet and outlet ports in a standard benchtop embryo incubator (Cook, Aus) at 37 °C in humidified 5% CO_2_, 5% O_2_, 90% N_2_. The microfluidic devices were placed inside 60 mm culture dishes and surrounded with embryo tested sterile water. To load embryos, media was then drawn through from the channel outlet until all embryos entered the central chamber (Supplementary material, Video 1 Loading and Video 2 Retrieval). 10 μl drops of pre-equilibrated KSOM were then added to channel inlet and outlet before culture at 37 °C under 5% CO_2_, 5% O_2_ in humidified nitrogen.

A total of 46 devices were used for embryo culture in experiments 1 and 2 and additional 30 for RT-PCR and metabolomics analysis.

### *In vitro* embryo culture in microdrops

Cryopreserved embryos (strain C57BL/6N) for Experiment 1 and 2 were supplied at zygote and 2-cell stages by MRC Harwell, UK. 1-cell murine embryos (B6C3F1xB6D2F1 strain, EmbryoTech, USA) were used for experiment 3 and 4. On the day of culture, murine presumptive zygotes were thawed following MRC protocols. Briefly, embryo straws were held in air for 30 s, then plunged into room temperature water until the contents had visibly thawed (around 10 s). The straws were cut at the seal and the plug bisected before pushing the contents into a 60 mm IVF hydrophobic culture dish. Embryos were incubated for 5 min before two 5 min washes in 100 μl M2 medium at 37 °C. Embryos were then washed through three 500 μl wells of pre-equilibrated KSOMaa before transfer to devices or culture microdrops. Culture microdrops were 1 μl/embryo in 35 mm hydrophobic IVF certified dishes (Nunc), covered with 5 mL of BioUltra mineral oil from Sigma Aldrich.

### Experimental design 1-Cell count, outgrowth assay

Cell counts were performed using the method described by Thouas et al. [45]. Briefly zona-intact blastocysts were first incubated in 500 μL of DPBS with 1% Triton X-100 and 100 μg/mL propidium iodide for 5s resulting in a labelled trophectoderm with red fluorescent signal. Blastocysts were then immediately transferred into 500 μL of fixative solution (100% ethanol with 25 μg/ml Hoechst 33258) and stored at 4°C overnight. Fixed and stained blastocysts were then mounted onto a glass microscope slide and gently flattened with a coverslip to facilitate the individual identification and counting of fluorescent cells. Fluorescent images were taken on a Zeiss AX1 Epifluorescence microscope and analysed in ImageJ.

Outgrowth assays were performed as described by Hannan et al. [86]. Blastocysts were imaged and measured using a Nikon ICSI microscope with RI viewer software.

### Experimental design 2 – Blastocyst rates, energy substrate consumption and mitochondrial polarization

To define the loading capacity of device culture, groups of 10, 20, 30 and 40 2-cell mouse embryos were cultured to the blastocyst stage in microfluidic devices in parallel to controls (10 embryos in one 10 μL culture microdrop) before metabolic profiling.

A total of 220 blastocysts were analysed in each treatment group from 5 different replicates. Glucose and pyruvate consumption were measured using spent media from devices or microdrops by the method of Guerif et al. [41] and expressed as mean pmol/embryo/hr ± SEM. Briefly, 10 μl reaction mixture was added to the base of a black 384 well microplate. 1μl spent media from device or microdrops was added to each reaction mixture well and the difference in NAD, NADH or NADP signal, measured at 340/460nm, was calculated. Concentration was calculated using a 6-point standard curve and expressed in terms of pmol/embryo/hr.

Day 8 blastocysts were labelled and imaged within devices using the ratiometric mitochondrial dye JC-10 (Molecular Probes). Briefly, a 1mg ml^−1^ JC-10 stock solution was prepared in KSOM culture media. JC-10 stock was diluted to 10ug ml^−1^ in pre-equilibrated M2 media (Millipore). Following removal of spent media for energy substrate assays, media was replaced with this JC-10 solution and incubated at 37 °C for 30 min. Control embryos were imaged on glass slides, while device embryos were imaged within devices to avoid stress. Imaging was performed on a LSM 780 confocal microscope, while image analysis was carried out in ImageJ. Data was represented as red fluorescence/total red+green fluorescence to account for any non-specific fluorescence across both channels. Higher values indicate a higher level of mitochondrial polarisation throughout the measured region of interest.

### Experimental design 3-Real-time PCR (qPCR) of single blastocysts

Groups of 10, 1-cell murine embryos were cultured in KSOM medium in 10 μL microdrops (control) and inside the devices. 90 blastocysts were analysed in each treatment group in 3 different replicates. Individual stage matched expanded blastocysts were recovered from devices or control microdrops, immediately transferred into 2 μl lysis buffer and frozen at −80 °C. For the construction of cDNA libraries of individual blastocysts we modified an existing protocol [87]. In detail, total RNA from single blastocysts was isolated using an RNAGEM-extraction reagent mastermix (RNAGEM Tissue Plus^®^, MicroGem International PLC, Southampton, UK). The total RNA was reverse-transcribed to cDNA using a first strand cDNA synthesis kit (Thermo Fisher Scientific Inc., UK). Quantitative PCR (qPCR) was performed on 6-fold diluted sample cDNA to analyze expression of 12 selected genes associated with blastocyst development and cell differentiation. Accession number, primer sequence and product length of target genes are presented in Table 2. Ten individual blastocysts were analyzed in each experimental group. mRNA expression was examined using SYBR Green Master PCR mastermix (Thermo Fisher Scientific Inc., UK) with an ABI 7500 RT-PCR System (Applied Biosystems). Data were analyzed with 7500 Software using relative quantification analysis.

**Table 2.**
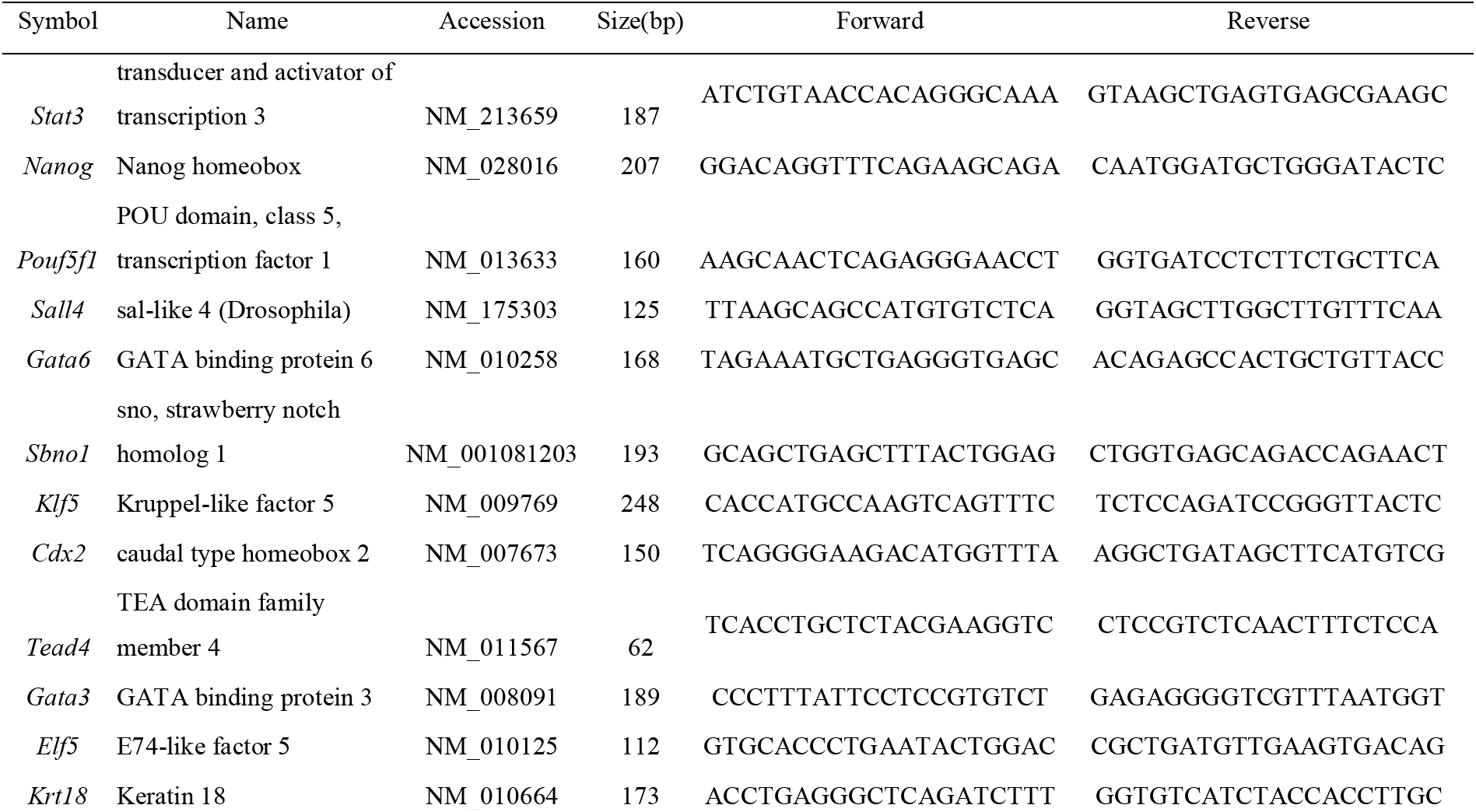
Summary of gene symbol, accession number, product length and primer sequences of the target genes.

### Experiment design 4 - Global untargeted metabolomics

To investigate if PDMS releases low molecular weight species or sequesters hydrophobic biomolecules from culture media [88], we analysed and compared samples of spent media (KSOM) collected from devices to samples of spent media collected from control KSOM microdrops using global untargeted metabolomics. 1-c murine embryos were cultured using the same embryo-to-volume ratio in microdrops or PDMS devices. In particular, 10 embryos were cultured in 40 μl KSOM drops and 10 embryos were cultured in each microfluidic device, where 20 μl drops were added to each inlet and outlet ports. Samples were collected from microdrops and microfluidic devices when embryos developed to fully expanded blastocyst stage, to allow stage-matched comparison of embryo metabolite production/consumption. Specific to this study, the blastocyst stage was achieved at day 4 in microdrops and day 5 in devices. Samples of spent media (40 μl) were collected from devices or microdrops to assess pre-implantation embryo metabolomics. As a control, spent media from devices without embryos was compared to media collected from microdrops without embryos using the same incubation time performed for embryo culture. These experiments were performed to investigate PDMS leaching and/or molecule absorption and adsorption. Culture media was frozen and stored at −80°C prior to sample preparation. Culture media samples (100 μl) were thawed on ice and prepared using previously described methods. Briefly, 300 μL of dry ice cooled methanol was added to individual culture medium samples and incubated overnight at −80°C; individual samples were spun down to remove proteins and subsequent supernatant used for analyses. Samples were separated and analysed using reverse-phase liquid chromatography connected to a a Thermo Scientific Q Exactive HF (LC-Hybrid Quadrupole-Orbitrap MS/MS) instrument using positive ion mode MS [89–91]. MS raw data were imported, processed, normalized, and reviewed using Progenesis QI v.2.1 (Non-linear Dynamics, Newcastle, UK). Resulting MS data was utilized for relative quantitation. The full collection of raw data has been published on Metabolomics Workbench [92]. Tentative and putative annotations [93] were determined using accurate mass measurements (<5 ppm error), isotope distribution similarity, and manual assessment of fragmentation spectrum matching from the Human Metabolome Database (HMDB), [94] Metlin, [95] MassBank, [96] and the National Institute of Standards and Technology (NIST) database [97]. Increased confidence in the annotation of many features was achieved by manually assessing spectral match and RT consistencies between experimental data and chemical standards within a curated in-house library.

### Chemicals

All chemicals were purchased from Sigma Aldrich (St Louis, MO, USA) unless specified otherwise.

### Statistical analysis

Data were analysed using GraphPad Prism 8 software. All data sets for first tested for fit to the normal distribution by D’Agostino-Pearson test for normality. In experiment 1, all normal data sets were compared by Student’s t-test, while all non-parametric data were compared by Mann-Whitney U test. In experiment 2, all data sets were non-parametric and therefore tested for significant differences between groups by the Kruskal-Wallis test with post-hoc Dunn’s test for multiple comparisons. In all instances, significance was determined as P<0.05.

For metabolic activity experiments, results were checked for statistical differences between groups by ANOVA with post-hoc Bonferroni test.

qPCR results were analysed by the comparative threshold cycle (Ct) method. Relative expression ratios were obtained using as internal control the mean of Ct values of eight housekeeping genes from the sample of interest. Student t-test statistics was used to compare gene expression levels between samples from the different groups.

For MS, compounds with <20% coefficient of variance (%CV) were retained for further analysis. Within Progenesis QI, a one-way analysis of variance (ANOVA) test was used to assess significance between groups and returned a P-value for each feature (retention time_m/z descriptor), with a nominal P-value ≤ 0.05 required for significance. Significant features were further filtered using a fold change threshold ≥ |2| deemed as significant.

## Supporting information

Video 1 Loading and Video 2 Retrieval

## Acknowledgments

We would like to thank Mr. F. Colucci for lab work and design contribution in the first experimental phase of the project, and Miss C. Espinilla Aguilar that completed the loading capacity experiment and analysis of polarisation ratio.

This project has been sponsored by the Medical Research Council Harwell Institute through the NC3Rs CRACK IT EASE Challenge (CRACKITEA-SP-3). It has been partially funded by the MRC CiC5 - Confidence in Concept 2018/19 (MC_PC_17165). This project has received funding from the European Union’s Horizon 2020 research and innovation programme under the Marie Skłodowska-Curie grant agreement No 748903.

## Author Contribution

V. Mancini participated to the design and optimization of the embryo culture device, design and completed the PCR analysis and generated the samples for MS analysis. Vanessa is now in a new position at the Department of Anatomy & Embryology, Leiden University Medical Center, Einthovenweg 20, 2333 ZC, Leiden, The Netherlands.

P. McKeegan carried out the experimental validation of the device, organized the embryo culture work and completed the analysis of blastocyst rate, outgrowth, mitochondrial polarisation ratio and GPL.

A.C. Rutledge performed metabolomics data processing and management of annotation activities; provided supervision and applied statistical techniques to study data. She also critically reviewed, and provided commentary and revision to the manuscript.

S.G. Codreanu designed the LC-MS/MS analytical method; processed the samples for metabolomics analysis and collected the MS data. She verified the overall quality and reproducibility of the results; provided revisions to the final manuscript.

S.D. Sherrod defined the best analytical method for the untargeted metabolomics analysis using the effluent from the microfluidic system. She mentored the core team on the metabolomics research activity, planning and execution as well as critically reviewed, and provided commentary and revision to the final manuscript.

J.A. McLean provided all the resources available for the metabolomic studies, including: reagents, materials, instrumentation, and computational resources for the metabolomics analysis. He also provided commentary and revision to the final manuscript.

H.M Picton defined the specifics for design and use of the microfluidic systems and advised on embryo culture and embryo quality assessment and grading. Analysed the IVF market, the expectations and needs in the field and provided guideline for the definition of a clinically relevant device.

V. Pensabene conceived the device design and characteristics and promoted its use in animal and human IVF. She defined the experimental protocols and the analytical methods, defined the full manuscript structure, provided guidance to the co-authors for the different activities and wrote introduction and conclusions.

## Additional Information

Supplementary information

Video 1 Loading

Video 2 Retrieval

## Competing interest

The Authors do not have any conflict of interest with the described research results.

## Notes

### Competing Interest Statement

The authors have declared no competing interest.

