## Supplementary material for "A new microfluidic concept for successful *in vitro* culture of mouse embryos": Video 1 Loading and Video 2 Retrieval

### Slide 1
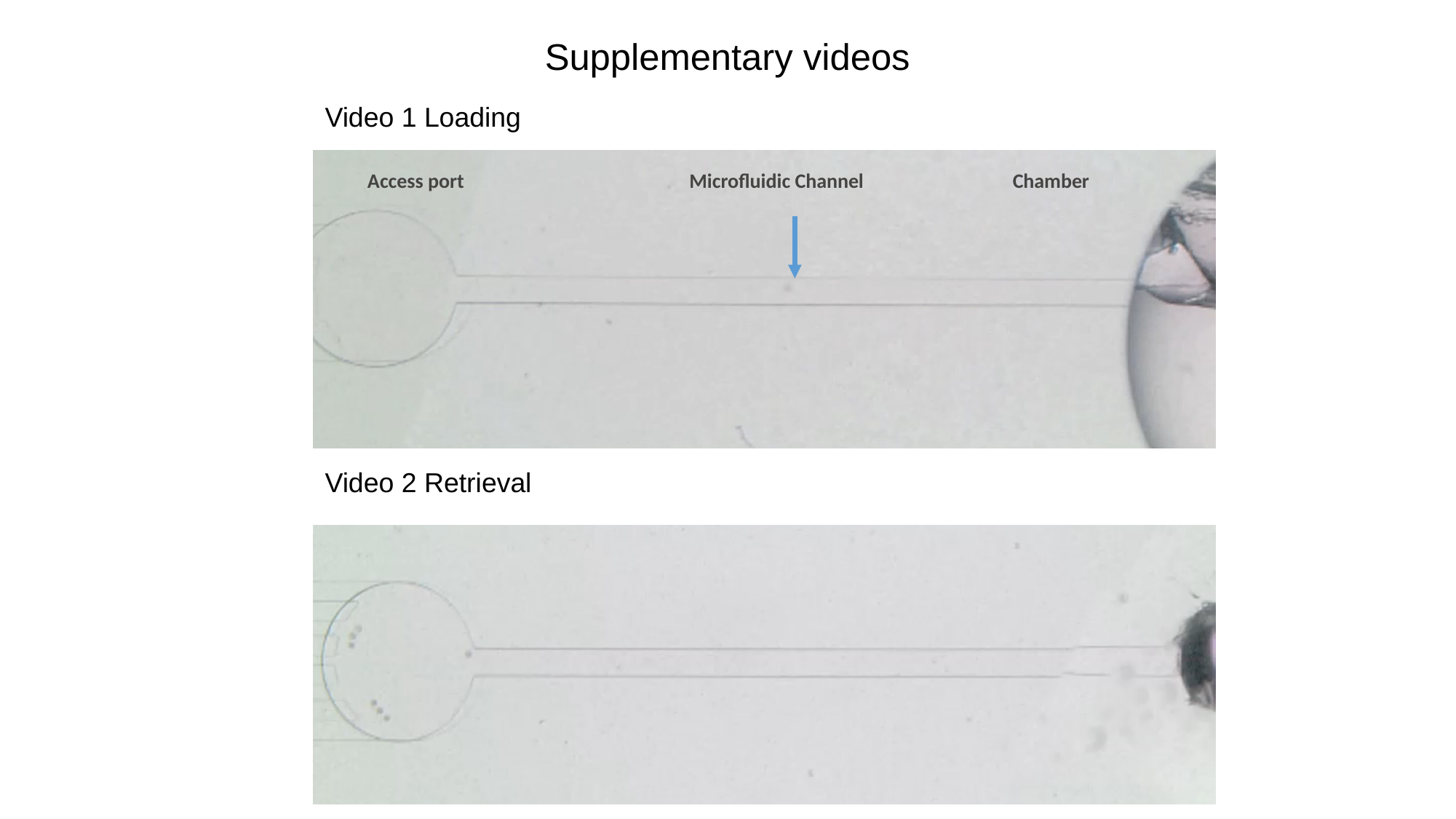

#### Supplementary videos
Video 1 Loading
Chamber
Microfluidic Channel
Access port
Video 2 Retrieval
