## Supplementary material for "A new microfluidic concept for successful *in vitro* culture of mouse embryos"

### Figure S1

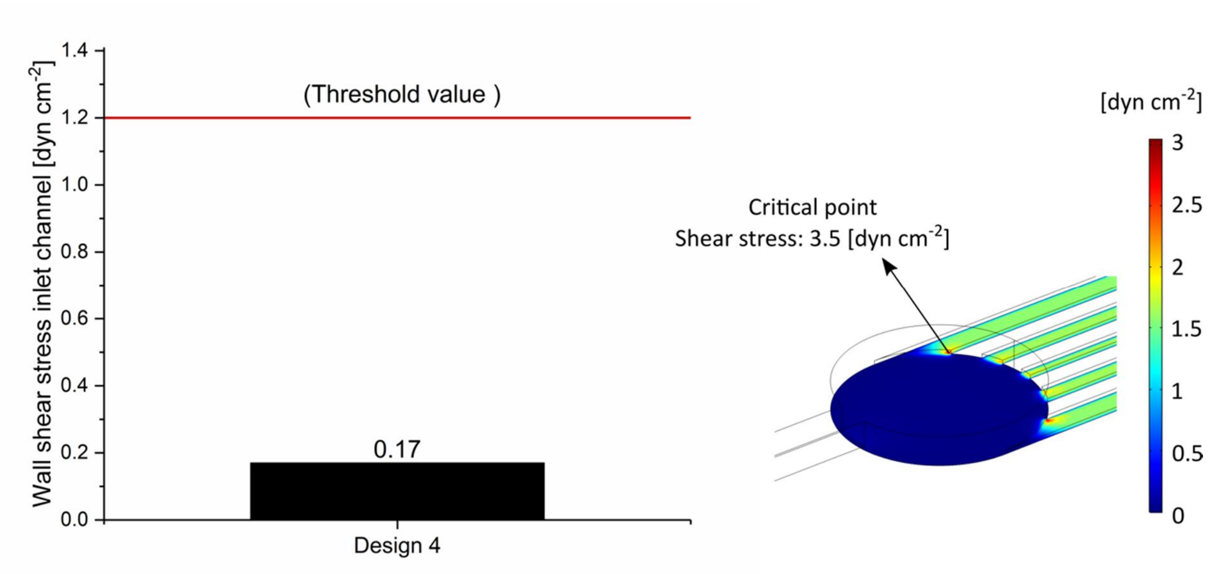


**Figure S1**. a) Computationally estimated inlet channels wall shear stress for the microfluidic device in its final version (Design 4), and b) fluid flow computational model of shear stress field surface plot during manual loading. The colour spectrum bar shows the shear stress field generated in the fluid systems.

***Figure S2***

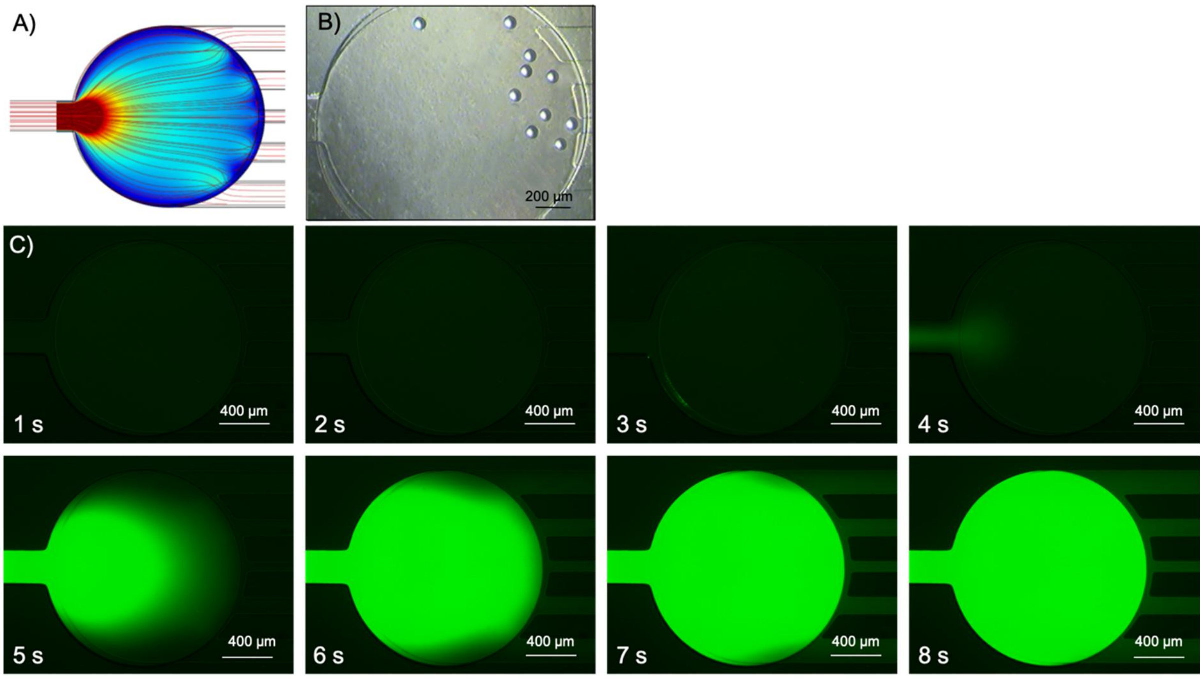


**Figure S2**. Real flow characterization within the culture chamber. A) Velocity magnitude surface and stream lines plot. B) Polystyrene beads spread within the culture chamber. C) Microfluidic device filled with a 0.05 mg ml$^{-1}$ fluorescein solution.

*Figure S3*

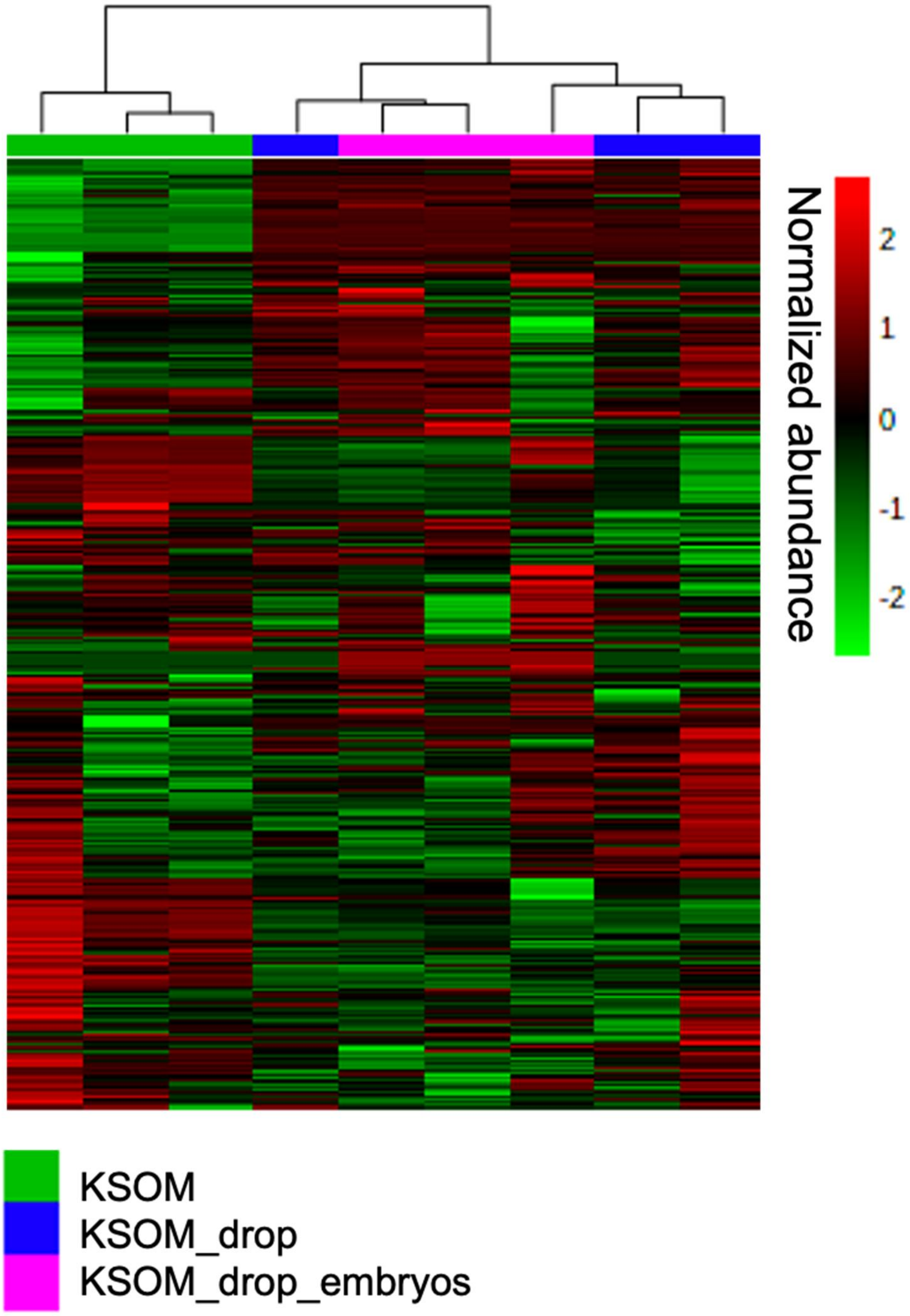


**Figure S3**. Heatmap analysis of media samples collected from control microdrop with and without embryos. Sample replicates are visualized in columns column based on hierarchical clustering, with metabolites presented on individual rows. Species are colored based on normalized abundance from red (high) to green (low).
